# Bacterial multispecies invasion of human epithelial bladder cells

**DOI:** 10.1101/2023.02.16.528894

**Authors:** Charlotte Abell-King, Alaska Pokhrel, Scott A. Rice, Iain G. Duggin, Bill Söderström

## Abstract

Urinary tract infections (UTI) are one of the most common bacterial infections worldwide. While the overall infection course is known on a macroscale, bacterial behavior is not fully understood at the cellular level and bacterial pathophysiology during multispecies infection is not well characterized. Here we establish co-infection models combined with high resolution imaging to compare single- and multi-species bladder cell invasion events in three common uropathogens: uropathogenic *Escherichia coli* (UPEC), *Klebsiella pneumoniae* and *Enterococcus faecalis*. While all three species invaded the bladder cells, under flow conditions the Gram-positive *E. faecalis* was significantly less invasive compared to the Gram-negative UPEC and *K. pneumoniae*. When introduced simultaneously during an infection experiment, all three bacterial species sometimes invaded the same bladder cell, at differing frequencies suggesting complex interactions between bacterial species and bladder cells. Inside host cells, we observed encasement of *E. faecalis* colonies specifically by UPEC. During subsequent dispersal from the host cells, only the Gram-negative bacteria underwent infection-related filamentation (IRF). Taken together, our data suggest that bacterial multispecies invasions of single bladder cells are frequent and support earlier studies showing intraspecies cooperation on a biochemical level during UTI.

## Introduction

Urinary tract infections (UTI) are amongst the most common bacterial infections globally ^1^. With an estimated 150 million people experiencing an UTI annually ^2^. Clinically, UTI are predominantly attributed to only one species of bacteria at a time, with Uropathogenic *Escherichia coli* (UPEC) being the most commonly diagnosed infectious agent at a reported prevalence between 70 and 95% ^3-7^. However, it is well established that there are a multitude of different bacterial species present in the bladder environment at any given time ^8^, and it is known that in acute UTI, polymicrobial infections are common, with both Gram-negative and Gram-positive pathogens present ^9,10^. Curiously, multispecies UTI are clinically underdiagnosed, especially if they contain Gram-positive species (*e.g.*, enterococci), and are often disregarded as sample contamination ^11^. It was found that in UTIs from 80 female patients with cystitis, that UPEC was over twenty-fold more common than *Enterococcus faecalis* (Gram-positive cocci) as the causative agent of the infections^12^. *E. faecalis* and UPEC are often associated with each other during outgrowth of cultures from UTI ^13,14^ and *E. faecalis* can suppress the immune activation, promoting UPEC virulence ^15^. Recent studies have investigated co-colonization in UTI models ^16,17^, but very limited information regarding bacterial invasion behaviours under liquid flow conditions simulating the bladder environment is available at high resolution at a single cell level.

Here we use two parallel approaches (‘flow’ and ‘dish’) to investigate differences in human epithelial bladder cell invasion rates of one, two or more bacterial species in the same infection experiment. The first approach is based on a flow channel model with constant exchange of nutrients ^18^, while the second approach use glass bottom petri dishes under constant orbital agitation. Pathogens used in this study were Gram-negative Uropathogenic *Escherichia coli* (UPEC) and *Klebsiella pneumoniae*, as well as the Gram-positive *Enterococcus faecalis,* as they are the three most commonly found organisms in healthcare-associated UTIs (HAUTIs) ^19^, and are commonly co-isolated in clinical settings ^20^, especially in samples from catheter-associated urinary tract infections ^21-24^. Challenging the human epithelial bladder cells with various combinations of the bacterial species and examining the system using high-resolution live cell fluorescence microscopy we show, that more than one species of bacteria frequently invades the same bladder cell.

## Results

### Flow and semi-static agitation affect invasion of human epithelial bladder cells by Gram-positive bacteria differently from Gram-negative bacteria

Initially, we established monospecies infections simulating the invasion phase of uncomplicated UTI using PD07i immortalized human epithelial bladder cells ^25,26^, challenged with either UPEC (strain UTI89 expressing cytoplasmic mOrange2), *K. pneumoniae* (strain TOP52 expressing cytoplasmic mCerulean3) or *E. faecalis* (strain SD234 expressing cytoplasmic GFP), to ascertain whether they were capable of infecting host cells in a previously established *in-vitro* UTI flow-chamber model and to establish baseline infection rates ^18,27^. This model is based on a commercial flow-chamber system (IBIDI I^0.2^ *μ-*Slides connected to NewEra syringe pumps) where constant flow is applied throughout the infection cycle to mimic bladder flow. In parallel, we also used a semi-static infection model using 35 mm glass-bottom Petri dishes under constant orbital agitation (50 rpm). In both models, the bladder cells were initially exposed to a total of ∼ 10^7^ bacterial cells of each species. All three pathogens invaded the epithelial bladder cells both in the flow-chamber and Petri-dish conditions (Fig. 1a-d).

**Fig. 1.**
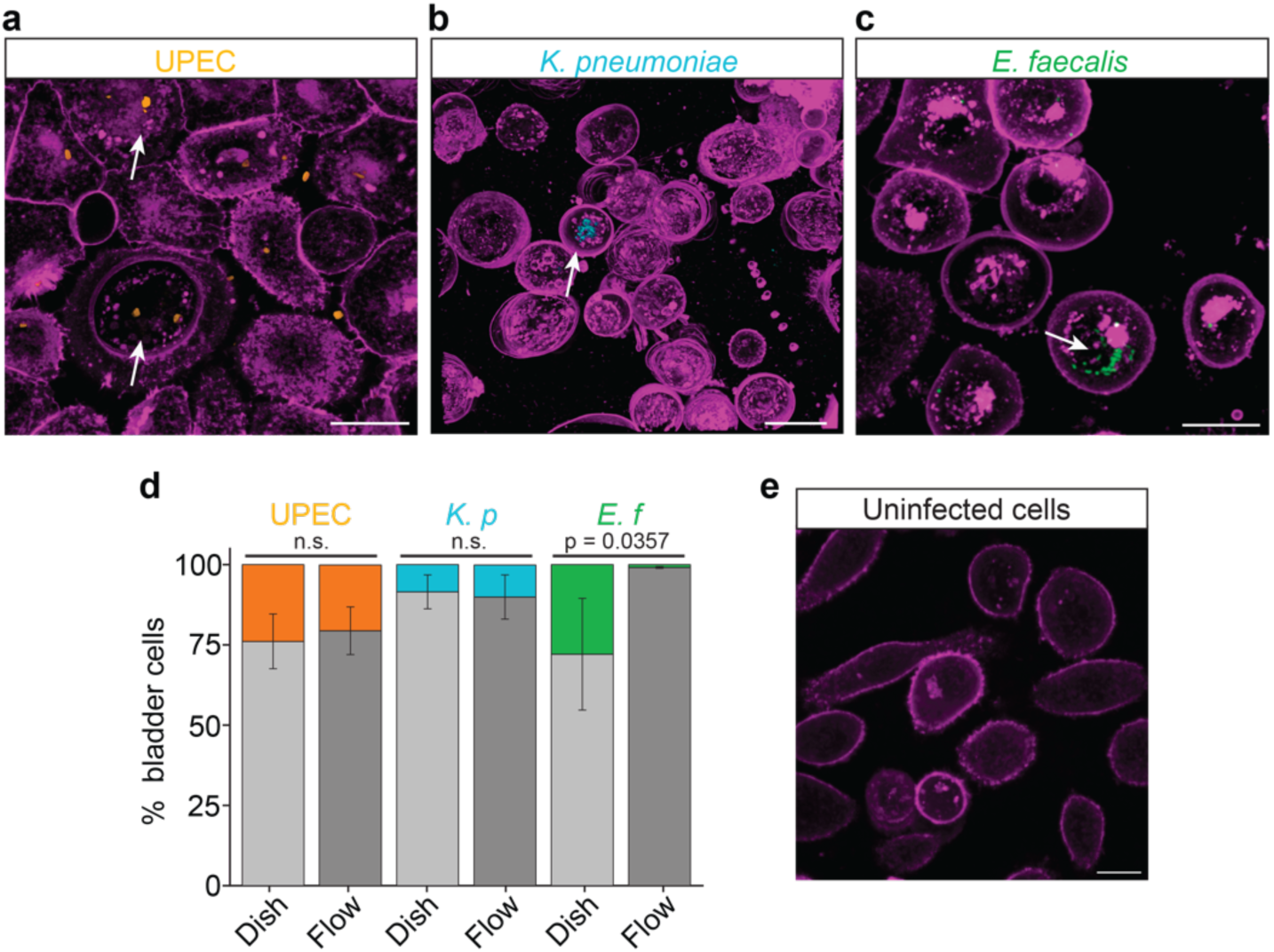
Single-species invasion of human epithelial bladder cells using Gram-negative and Gram-positive uropathogens. *Immortalized epithelial bladder cells PD07i (cytoplasmic membranes labelled with CellBright650, magenta) grown on a 35-mm glass bottom petri-dish or in a flow chamber were infected by either **a**, UPEC (strain UTI89 expressing cytoplasmic mOrange2); **b**,* K. pneumoniae *(strain TOP52 expressing cytoplasmic mCerulean3); or **c**,* E. faecalis *(strain SD234 expressing cytoplasmic GFP). White arrows indicate internalised bacteria. **d**, Proportion of bladder cells infected (colour) and uninfected (grey) in various conditions and bacterial species. Graphs are based on values from 3 infections for each condition. n.s. = not statistically significant. p < 0.05 = statistically significant*. *K. p =* K. pneumoniae*. E. f =* E. faecalis*. Dish = 35 mm glass bottom petri-dish. Flow = 15 μl min^-1^ in flow channel (IBIDI μ-Slide I^0.2^). **e**, PD07i cells maintained without bacteria. Images are from petri-dish experiments. Scale bars = 20 μm. Note that the bottom layers containing only epithelial membrane of the Z-stacks were omitted for better visualization of internal bacteria*.

At 20 h post inoculation, UPEC had invaded the most bladder cells of the tested Gram-negative species (23.9 ± 8.5 % of the bladder cells in the semi-static petri dish model and 20.6 ± 7.4% in the flow model, Mean ± S.D., n > 432 from three independent infection experiments), while *K. pneumoniae* invaded approximately half of this number (8.49 ± 5.3% in the dish model and 10 ± 6.9 % in the flow model, n > 321 from three independent infections of each condition) (Fig. 1d). *E. faecalis* showed the highest overall invasion with an average 27.9 ± 17.4% of the bladder cells invaded in the dish model (n = 335, from 3 experiments) (Fig. 1d). In contrast, *E. faecalis* invaded less than 1% of the bladder cells under the flow-chamber conditions (n = 462, from 3 independent experiments). For UPEC and *K. pneumoniae* the variations between the models were statistically nonsignificant, while for *E. faecalis* they were p = 0.0357 (statistically significant).

### Elevated invasion rates for UPEC in dual species infection with E. faecalis

Since so few bladder cells were invaded by *E. faecalis* in the flow-chamber model, the culture-dish model was mainly used to follow invasion during multispecies UTI. We co-inoculated the bladder cells with equal numbers of UPEC and *E. faecalis* (a combined total of ∼ 10^7^ bacteria ^15^). Based on images of more than 560 randomly inspected bladder cells at ∼ 20h post inoculation (regions of interest [ROIs] were chosen based on fluorescence signal from bladder cell membranes, and Z-stacks were acquired for each ROI) from three independent experiments, almost half (∼ 45%, Fig. 2a - b) were invaded by at least one type of bacterial species. Most infected bladder cells were only invaded by one species (Fig. 2a, arrows).

**Fig. 2.**
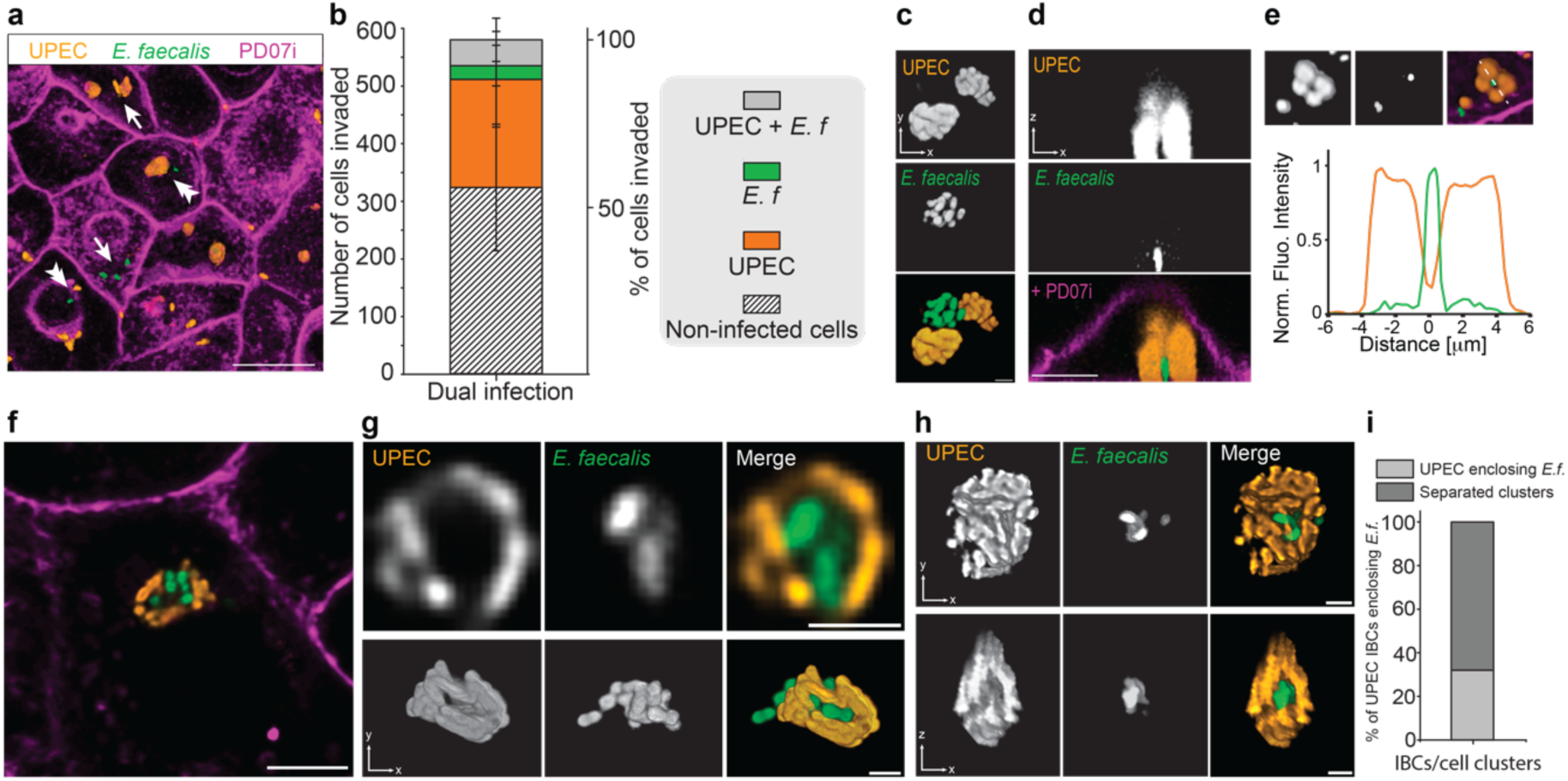
Bladder cell invasion during UPEC and E. faecalis co-infections. ***a****, PD07i (membranes labelled with CellBright 405 or 650) cells co-infected with* UPEC *(orange) and* E. faecalis *(green). Scale bar 20 = μm. **b**, ratio of invaded PD07i cells. E. f = E. faecalis **c**, Typical intracellular* UPEC *(orange) and* E. faecalis *(green) clusters.* UPEC *appeared more tightly packed than* E. faecalis*. Scale bar 1 μm. **d** and **e**, Low resolution Z-stacks showing UPEC surrounding* E. faecalis *in co-invaded PD07i cells. Scale bar **d** = 5 μm. The graph in **e** shows fluorescence intensity traces for the white dash line in the image above. UPEC (orange) and* E. faecalis *(green). Dotted scale line = 12 μm. **f**, **g** and **h**, Representative single images and confocal 3D reconstructions of* E. faecalis *(green) surrounded by UPEC (orange). Scale bar **f** = 10 μm. Scale bars **g**, **h** = 2 μm. **i**, Percentage of dual invaded PD07i cells with UPEC cells enclosing* E. faecalis cells *(as shown in **g** and **h**). Note various orientations of reconstructed Z-stacks*.

In these dual-species infections, UPEC alone were internalised in 33% of cells (compared to ∼ 24 % in single species dish infections [Fig. 1d]), whereas *E. faecalis* alone were internalised in only 4% (Fig. 2b) (compared to ∼ 28% in single species dish infections [Fig. 1d]). The percentage of bladder cell invasions of either or both bacteria in dual-species infections was almost 45% (43.6 ± 19.3% (n = 564 cells, from 3 experiments)). Surprisingly to us, a relatively high number, 44 of 564 or approximately 8% of the individual bladder cells were invaded by both bacterial species at the same time (Fig. 2a, arrow heads).

We noticed that the arrangement of the internalised bacteria was distinct for each species; UPEC formed condensed IBCs as previously established ^28^, while *E. faecalis* were often organized in more loosely organised clusters in which single cells were readily resolved (Fig. 2c). Initial 3D reconstructions of low-resolution images suggested that UPEC clusters were often in close spatial proximity of the *E. faecalis* cells (Fig. 2d - e). With increased resolution, it became evident that tightly packed UPEC cells formed multicellular communities that frequently surrounded one or a few *E. faecalis* cells (Fig. 2f - h). This type of UPEC ‘encirculation’ of *E. faecalis* cells was apparent in a substantial fraction of the total observed UPEC IBCs (32%) during dual species invasion (Fig. 2i). We speculate that *E. faecalis* and *E. coli* might cooperate to increase the likelihood of prolonged infection ^15^.

### Lower overall infection rates of bacteria in dual and triple-species infections when K. pneumoniae is present

To investigate whether other mixed bacterial cultures would co-invade PD07i cells and exhibit similar interactions as seen for UPEC / *E. faecalis*, we carried out infection experiments with other combinations of bacteria: *E. faecalis* and *K. pneumoniae* (Gram-positive/Gram-negative pair), UPEC and *K. pneumoniae* (Gram-negative/Gram-negative pair) or all three species (UPEC, *E. faecalis* and *K. pneumoniae*). At 20h post inoculation, we detected host cells containing all combinations of the bacteria investigated (Fig. 3a – c), but the invasion frequencies varied substantially.

**Fig. 3.**
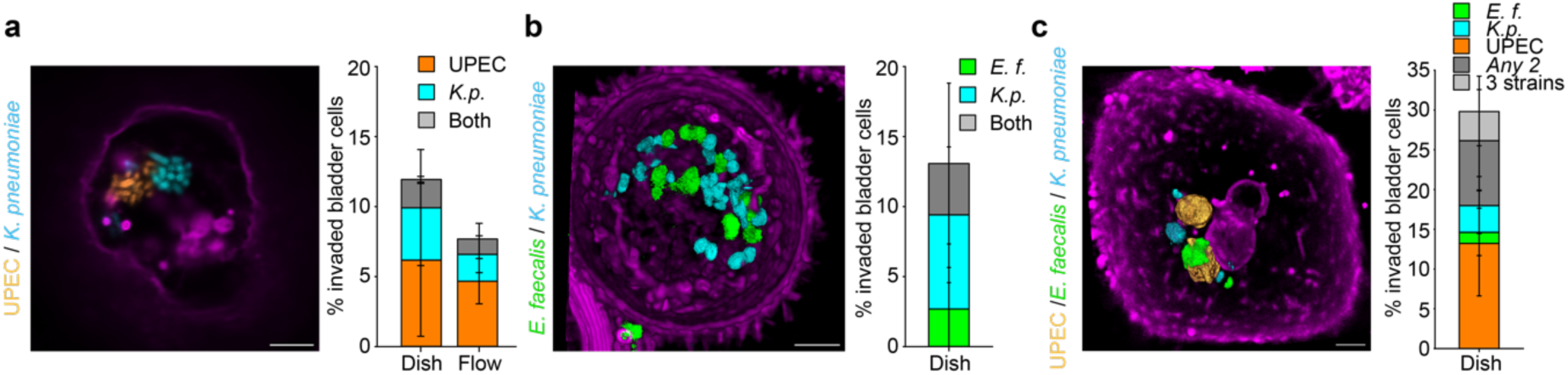
Prevalence of bacterial invasion in dual- and triple-species infections. *Representative images of PD07i cells invaded by **a**,* UPEC *and* K. pneumoniae *(2D confocal image); % of invasion (n = 519), Dish:* UPEC *6.1 ± 5.4 % (mean ± S.D.), K.p. 3.7 ± 4.2 %, both 2 ± 2.2 %. Flow:* UPEC *4.6 ± 2.6 %, K.p. 1.9 ± 1.8 %, both 1.35 ± 1.9 %. **b**,* E. faecalis *and* K. pneumoniae *(deconvolved 3D-reconstruction of a confocal Z-stack, n = 463). % of invasion: E.f 2.7 ± 3 %, K.p. 6.8 ± 4.9 %, both 3.7 ± 5.8 %. **c**,* UPEC, E. faecalis *and* K. pneumoniae *(deconvolved 3D-reconstruction of a confocal Z-stack, n = 349). % of invasion: UPEC 13.3 ± 6.7 %, E.f 1.3 ± 3 %, K.p. 3.4 ± 3.6 %, Any 2 species 8.2 ± 6.4 %, all 3 strains 3.7 ± 4.4 %. PD07i membranes labelled with CellBright 650. Scale bars = 4 μm. E. f =* E. faecalis*. K. p* = K. pneumoniae*. Note, different scales on y-axis*.

In contrast to the UPEC / *E. faecalis* case, encirculation was not observed when *E. faecalis* was co-infected with *K. pneumoniae* (4 independent infection experiments, n = 463 cells in total), nor when the two Gram-negative species UPEC and *K. pneumoniae* were used in the same infection (4 independent infection experiments, n = 519 cells in total). UPEC and *K. pneumoniae* often formed dense IBCs adjacent to one another but did not encase or mix with each other (Fig. 3a). Overall, UPEC and *K. pneumoniae* invaded ∼ 12% of all bladder cells in the Dish model and ∼ 8% of all bladder cells in the flow channel model, however they coinfected only 1 - 2% of bladder cells both in either model (Fig. 3a).

During *E. faecalis* and *K. pneumoniae* co-infection, we noticed that while they were capable to co-invade bladder cells, we did not observe dense IBCs (Fig. 3b). *E. faecalis* has previously been shown to antagonize *K. pneumoniae* biofilm formation during mixed growth albeit under different conditions compared to the intracellular conditions in the present study ^29^. The total percentage of invaded bladder cells in a *E. faecalis* and *K. pneumoniae* dual species combination was ∼ 13%, with 3.7% of all observed of bladder cells invaded by both (Fig. 3b).

For three-species infections (UPEC, *E. faecalis* and *K. pneumoniae)*, UPEC / *E. faecalis* again showed similar organizational patterns as they did in two species infections, while *K. pneumoniae* was predominately localized separate from the other two (Fig. 3c). In the three species infections roughly 30% of bladder cells were invaded by at least one species, with almost 4% invaded by all three (Fig. 3c, n = 349, 3 independent infection experiments). This was less than the invasion frequency of the UPEC / *E. faecalis* dual infections where ∼ 45% of the bladder cells were invaded by at least one type of bacteria (Fig. 2b), suggesting that *K. pneumoniae* could have an antagonistic effect on invasion efficiency during multi-species infections.

### Gram-negative bacteria undergo infection-related filamentation in multispecies infections, while Gram-positive do not

During murine model UTI, UPEC have been seen to undergo morphological changes during intracellular bacterial community (IBC) formation and dispersal, where UPEC grow into long filamentous cells ^30^. These morphologies were later also seen in female patents with cystitis ^12^ and more recently also confirmed in flow-chamber models using human bladder cells and urine ^18,27,31,32^. To determine whether single and multi-species co-infections undergo the same morphological changes under the established conditions, we next visualized cells sampled after the dispersal stage of infection by exposing infections to human urine (20 h) ^18,27^. Both UPEC and *K. pneumoniae* formed filaments of several hundreds of microns long (Fig. 4a - b). For *K. pneumoniae*, similar morphology cycles have been previously observed in murine models ^33^, but not in *in-vitro* models using human epithelial bladder cells and urine. The average length of both UPEC and *K. pneumoniae* filaments from infections was ∼ 50 μm (Fig. 4d). By comparison, UPEC and *K. pneumoniae* rods grown in LB medium were approximately 3.5 μm (Fig. 4d). We classified cells as filaments if they were at least two times the average WT cell length, *i.e.*, 7 μm. We observed that the number of cells that underwent filamentation (but not the average length of filaments) was lower in the dish model compared to the flow-chamber model. While the molecular reasons for this is not currently clear, we speculate that proficiency of morphology changes may be connected to flow and shear-force dynamics, or the continual exchange of constituents in the medium ^31^. In contrast to the Gram-negative species, *E. faecalis* cells did not filament or grow significantly larger upon exposure to human urine (Fig. 4c, 4d). Average lengths after an infection were ∼ 1.4 μm (n = 215) from petri-dish infection and ∼ 1.6 μm (n = 102) from flow chambers which was similar to lengths for *E. faecalis* cells grown in BHI medium only (1.75 (n = 238)) (Fig. 4d). Similar results for filamentation were observed during triple-species infections (Fig. 4e). Taken together, these observations suggest that infection-related filamentation may be a broadly occurring phenomenon in uropathogenic Gram-negative bacteria and it also transpires during multispecies infections.

**Fig. 4.**
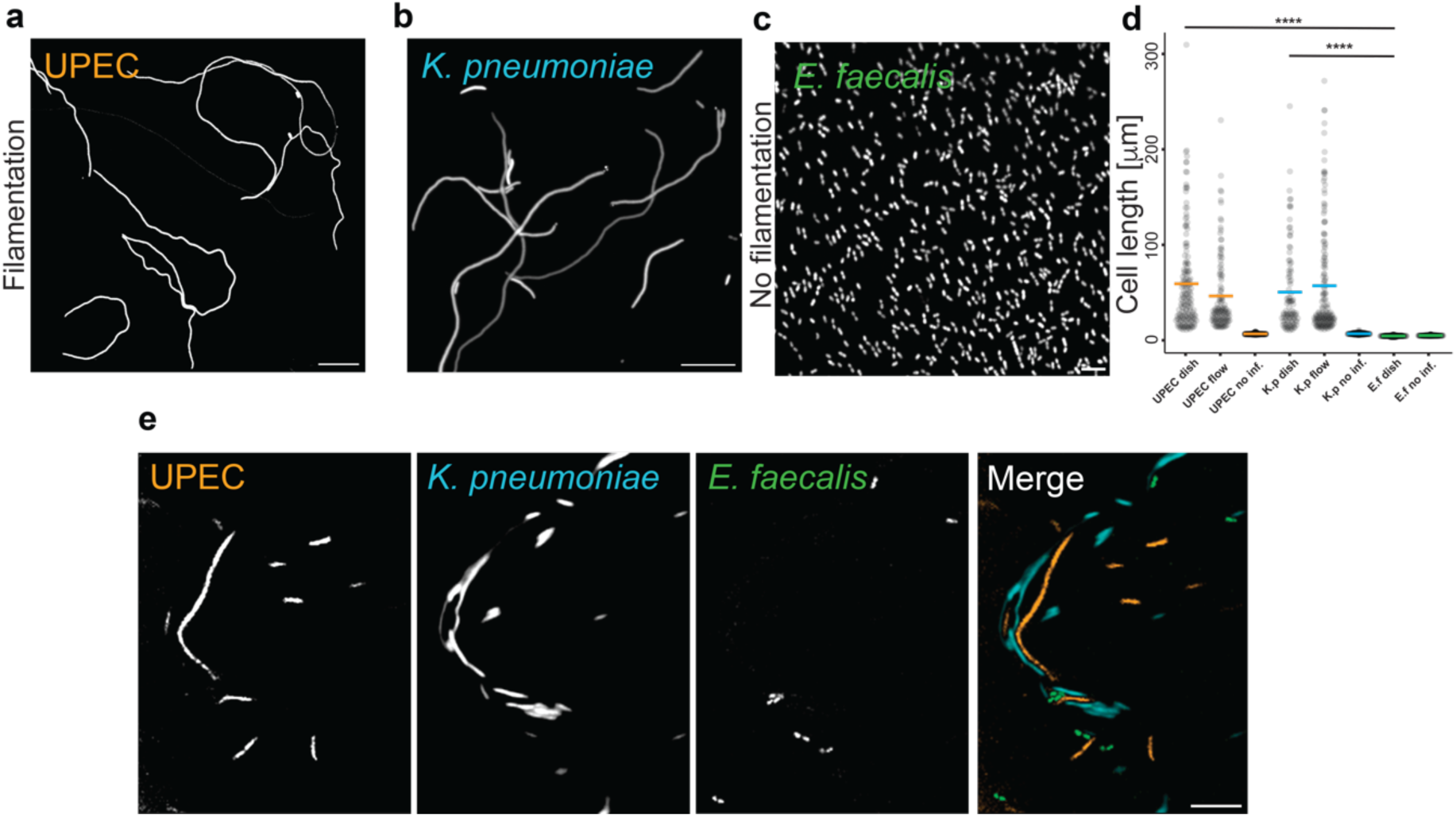
Gram-negative, but not Gram-positive, species undergo infection-related filamentation in both single and multi-species infections. *Representative images of bacteria after being exfoliated from PD07i cells during single species infections. **a**,* UPEC*, **b**,* K. pneumoniae, ***c***, E. faecalis*. Note all strains retain their fluorescence throughout invasion and exfoliation. **d**, Average lengths of bacterial filaments* (UPEC and K. pneumoniae*) and cells* (E. faecalis*) after 20 h of urine exposure during infections. The average length of* UPEC *filaments was 56.0 ± 59.9 μm (n = 158) in dishes, and 43.3 ± 40.05 μm (n = 122) in flow.* K. pneumoniae *filaments were 47.3 ± 45.3 μm (n = 98) in dishes and 54 ± 57.8 μm (n = 156) in flow and.* UPEC *and* K. pneumoniae *rods grown in LB media were on average 3.46 ± 0.69 μm (n = 187) and 3.62 ± 0.76 μm (n = 192), respectively. Average lengths of* E. faecalis *after an infection were 1.42 ± 0.4 μm (n = 215) from petri-dish infection and 1.59 ± 0.41 μm (n = 102) from flow chambers. Lengths for* E. faecalis *cells grown in BHI media only was 1.74 ± 0.31 μm (n = 238). Dish = 35-mm glass bottom petri-dish. Flow = 15 μl min^-1^ in flow chamber (IBIDI I^0.2^). **e**, Filamentation of* UPEC *and* K. pneumoniae *but not* E. faecalis *during a representative three species infections. Scale bars A, B and E = 10 μm, C = 4 μm*.

## Discussion

Numerous studies have investigated UTI microbial interactions and biofilm formation *in-vitro* in liquid media (*e.g.*, rich media, synthetic or human urine) and on catheters ^34-37^, their effect on immune activation^15,17,38,39^, and *in-vivo* in animal models^40-42^ (a list on further reading here: ^8^). However, hardly any studies have looked at bacterial co-invasion behaviour at single bladder cell level. Here, to study and better understand bacterial invasion during multispecies UTI on single cell levels in a systematic way, we established a set of multi-species UTI *in-vitro* models. We visually compared invasion frequencies of three common urinary tract pathogens, Uropathogenic *Escherichia coli* (UPEC), *K. pneumoniae* and *E. faecalis,* using human bladder cell *in-vitro* infection models under two different conditions (‘flow’ and ‘dish’). In the flow condition, UPEC showed highest overall invasion frequency, followed by *K. pneumoniae*, while *E. faecalis* hardly invaded any bladder cells. Differences in invasion frequency between UPEC and *K. pneumoniae* has been observed previously in murine models, and was attributed differences in adhesion capabilities through lower type 1 pilus expression of *K. pneumoniae* ^33^. The invasion frequencies of UPEC and *K. pneumoniae* were only marginally impacted by switching from continuous flow to circular agitation. On the other hand, the Gram-positive *E. faecalis* exhibited a large variation in invasion frequency between the models tested in this study. While *E. faecalis* is a well-established uropathogen ^43,44^, our observations suggest a weakened ability to adhere and invade bladder cells under the flow conditions in our model. At this point we do not fully understand the details resulting in the large variation in bladder cell invasion of *E. faecalis* between our flow and dish models, but similar behaviours have previously been linked to the reduced adherence capabilities of *E. faecalis* ^11,43,45,46^.

In multispecies infection experiments, we show that multiple species of bacteria can invade the same bladder cell. While molecular details governing intracellular coexistence of bacteria during UTIs remains elusive, our observations indicate that whenever UPEC and *E. faecalis* are introduced together the overall rate of infection increases. Similar cooperation and increase in pathogenicity by UPEC and *E. faecalis* have been suggested based on results from murine models ^15^. The observation that UPEC readily encirculate *E. faecalis* is perplexing and its biological significance is unclear but suggests a spatial importance of the previous observations that these two species cooperate on a biochemical level to increase pathogenesis ^40,42,47,48^.

In contrast, whenever *K. pneumoniae* was introduced into a multispecies infection situation was the average invasion frequency always lower than in the corresponding single species infections. Supporting this observation, *E. faecalis* has previously been implicated in suppressing *K. pneumoniae* growth during polymicrobial biofilm formation in human urine ^29^. This suppression by *E. faecalis* seems also be extended to other Gram-negative species (*i.e.*, *Pseudomonas aeruginosa*) ^49^.

Both infection models presented in this study caries its own merits, and each may be favoured depending on the research question in mind, *e.g.,* ‘Flow’ could represent bladder release, while ‘Dish’ would possibly depict a resting bladder situation. We believe that the observations presented in this study will initiate a spark for further investigation into detailed genetics studies exploring dedicated signalling pathways and potential quorum sensing in multispecies UTIs at a single cell level. This will undoubtably play an increasingly important role in understanding competing or cooperative behaviours of bacteria during infections as antimicrobial resistance continue to rise.

## Materials and Methods

### Human Ethics

This study was approved by the UTS Human Research ethics committee with approval numbers HRCH REF No. 2014000452 and HREC ETH22-7590. All urine doners gave their approval to participate in this study by informed consent under this study’s ethics approval numbers above.

### Plasmid construction

pCAK1 and pAP1 plasmids were constructed by replacing the GFP sequence in pGI5^18^ with mCerulean3 and mOrange2 sequences, respectively. The mCerulean3 and mOrange2 gene fragments were amplified from pEB1-mCerulean3 and mOrange2-pBAD plasmids, both obtained from Addgene (Addgene plasmids #103973 and #54531) ^50,51^. To construct pCAK1 and pAP1, pGI5 was digested with *Nco*I and *Bam*HI, and assembled with mCerulean3 and mOrange2 PCR products containing 20-30 bp homologous regions from pGI5 on both the ends. This was performed using a *in vivo* DNA assembly method ^52^. The final products were confirmed by Sanger sequencing and fluorescence microscopy.

### List of Primers

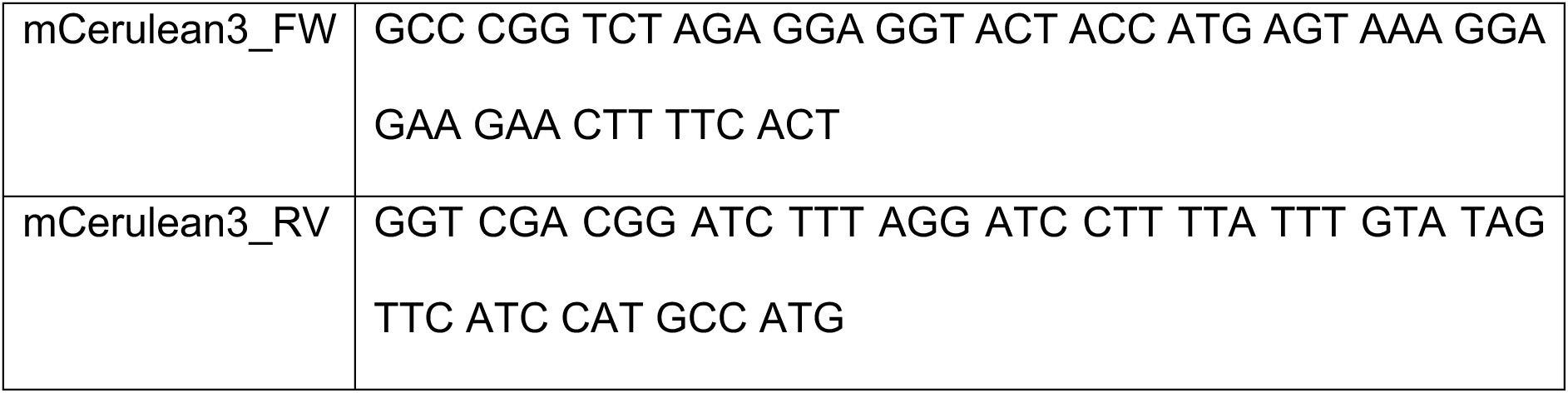
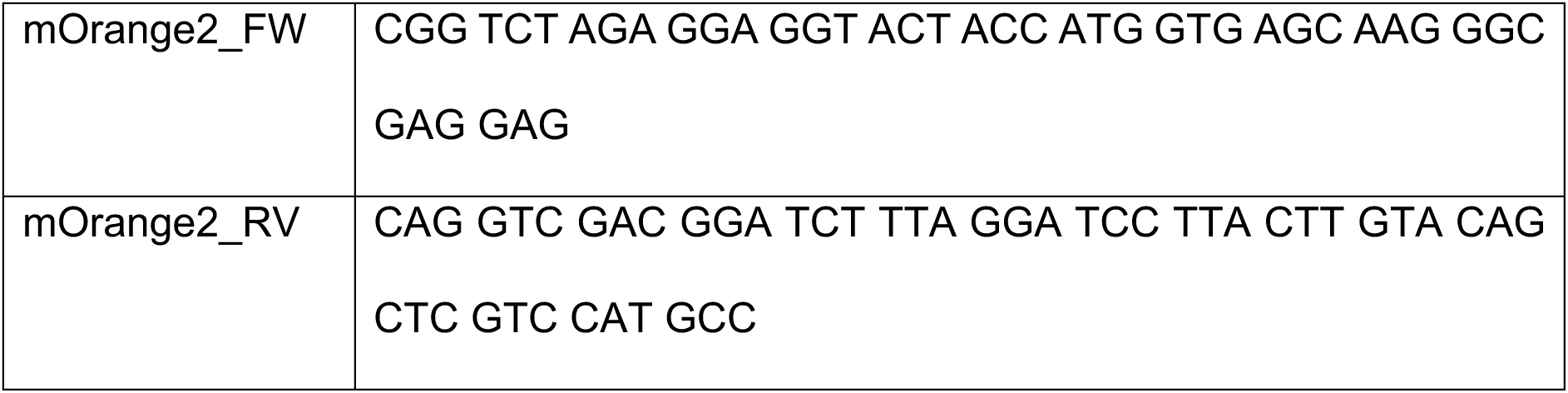

### Bacterial growth

Bacterial strains were used as follows: Uropathogenic *Escherichia coli* UTI89 transformed with mOrange2 (pAP1) or mCherry (pGI6) ^53^, *Klebsiella pneumoniae* TOP52 transformed with mCerulean3 (pCAK1) or sfGFP (pGI5), and *Enterococcus faecalis* OG1RF expressing GFP from the chromosome ^54^. All fluorescent proteins were produced as constitutively expressed freely diffusing molecules in the cytoplasms of the corresponding bacterial strain. A single colony of respective strain was grown overnight in a 20 ml Luria Broth (*E. coli* and *K. pneumoniae*) or Brain Heart Infusion (*E. faecalis*) culture at 37°C without shaking to induce expression of type-1 pili which facilitates adhesion to the bladder cells during infection. Antibiotics were added when appropriate (Spectinomycin 50 μg ml^-1^, Rifampicin 100 μg ml^-1^). The following morning, the cultures were pelleted and resuspended in 1 x PBS added to the infection model.

**Table 1:** Bacterial species used in this study.

| Species | Strain | Source | Fluorescent Protein | Plasmid Or Chromosome |
| --- | --- | --- | --- | --- |
| *E. faecalis* | OG1RF | Prof. Garsin (Uni Texas) | GFP | Chromosome |
| *UPEC* | UTI98 | Lab strain | mOrange2 | Plasmid (pAP1) -this study |
| *UPEC* | UTI98 | Lab strain | mCherry | Plasmid(pGI6)<sup>53</sup> |
| *K. pneumoniae* | TOP52 | Prof. Hultgren (Uni Washington) | mCerulean3 | Plasmid (pCAK1) -this study |
| *K. pneumoniae* | TOP52 | Prof. Hultgren (Uni Washington) | sfGFP | Plasmid(pGI5)<sup>18</sup> |

### Bladder cell growth

Immortalized Epithelial bladder cells (PD07i ^25,26^) were grown and maintained in EPILIFE media supplemented with Human Keratinocyte Growth supplement (HKGS) in 5% CO_2_ at 37 °C. Cells were split as required using standard Trypsinization methods upon reaching ∼ 80% confluency.

### Human urine preparation

Human urine was collected from both male and female donors. Samples were collected in the mornings and stored at 4 °C for at least 2 days before further processing. Samples were pelleted at 3000 rpm, the supernatant filtered through a 0.2 μm filter and aliquoted into 50 ml falcon tubes before placed in – 20 °C for storage until use. Urine was only used if the pH was between 5 - 6.5 and the Urine Specific Gravity (USG) was in the range between 1.024 to 1.030, values that has been shown to produce high degree of filamentation ^18,27^.

### Infection models

#### Flow model

The *in-vitro* UTI flow channel model has been described previously ^18,31^. Here only slight changes were introduced as follows, μ-Silde I^0.2^ Luer (IBIDI #80166, total channel volume 50 μl) flow channels were seeded with bladder cells according to the manufacturer’s recommendations. The channels were connected to a New Era pump system for continuous flow of nutrients, this was left to run until a confluent layer of bladder cells formed ^27^. Bacterial cells resuspended in PBS at an OD_600_ of 0.2 - 0.4 were introduced fully in the flow channels and flushed at a flow rate of 15 μl min^-1^ for 10-15 minutes (resulting in a total of ∼10^7^ bacterial cells run over the bladder cells depending on concentration and species, as determined by CFU counts), before fresh EPILIFE media again was flown over the cells for 7 hours (initial 10 minutes at 100 μl min^-1^ to flush out excess bacteria, then 15 μl min^-1^ for the reminder of the time) to allow for invasion. After this, the media was exchanged to EPILIFE containing Gentamycin (f.c. 100 μg ml^-118^, for the dual infection of UPEC and *K. pneumoniae* was on additional 100 μg ml^-1^ of Ampicillin added) for 13 hours to eliminate all extracellular bacteria. To monitor formation of intracellular bacterial communities, channels were taken to the microscope for imaging at this point. Immediately prior to imaging, bladder cell membranes were stained with CellBrite405 or 650 in PBS (f.c. 1:100, with the addition of CellBrite Enhancer to mask intracellular fluorescence according to the manufacturer’s recommendations) under flow (15 μl min^-1^) for 40 minutes, and washed twice with 1 x PBS. Channels were re-filled with EPILIFE media to sustain cell health during imaging.

To generate filaments, after subjecting samples to the EPILIFE/antibiotics mixture, human urine was added to the flow system (15 μl min^-1^) for 20 hours. Bacterial samples were collected through the back-end of the flow channels, washed once in 1 x PBS, resuspended in LB or BHI. 4 μl of respective culture was placed on 2 % agarose pads (w/w) and directly taken for imaging.

#### Dish model

In the semi-static infection model a similar experimental workflow was followed as for the flow model with the same time intervals in changing media, with the exception that all steps were carried out in a 35 mm Petri dish (IBIDI glass bottom dish with glass coverslip bottom (#1.5), pre-sterilized, cat number #81218-200) placed in a CO_2_ incubator at 37 °C with 50 rpm orbital agitation. For the dish models the total amount of bacteria (∼10^7^, roughly corresponding to an MOI of 100) was distributed evenly over the bladder cells at one time point only. Bacteria were incubated for 15 minutes before liquid was aspirated of and fresh EPILIFE media (2 ml) was added and grown for 7 hours. Following this, media was exchanged to EPILIFE containing Gentamycin (f.c. 100 μg ml^-118^), for 13 hours to eliminate all extracellular bacteria. To monitor formation of intracellular bacterial communities, channels were taken to the microscope for imaging. Immediately prior to imaging were bladder cell membranes stained with CellBrite405 or 650 in PBS (f.c. 1:100, with the addition of CellBrite Enhancer to mask intracellular fluorescence according to the manufacturer’s recommendations) for 40 minutes, washed twice in 1 x PBS and again covered with 2 ml EPILIFE to sustain cell health during imaging.

To generate filaments in the Petri dish model, 1 ml urine was added and manually exchanged once an hour for the first 4-5 hours before a final resuspension of 3 ml was done and left in the incubator for 15 hours.

### Imaging

To minimise biased imaging, regions of interest [ROIs] were chosen based on fluorescence signal from bladder cell membranes only, and Z-stacks were acquired for each ROI. Imaging was performed on a Leica Stellaris confocal microscope equipped with a 63x oil objective enclosed in an environmental chamber operated at 37 °C and 5 % CO_2_ (Oko-Lab). The fluorophores were excited by a white laser at optimized wavelengths, and emission was collected in pre-set system optimised detector intervals for AlexFluor405, mCerulean3, EGFP, mOrange2 and AlexaFluor647 depending on experiment, to minimize channel crosstalk. Z-stacks were always acquired to validate that the bacteria in fact were inside the bladder cells. Image size was either 2048 x 2048 or 4096 x 4096, with pixel size 90 and 45 nm, respectively. The pinhole was set to 1 AU and Z-stacks were collected with software optimized step length of either 125 or 250 nm (30-99 images per stack, depending on step length and thickness of the bladder cell in question). 3D reconstruction and deconvolution of Z-stacks were performed in the Leica LAS software and further visualised in FIJI (ImageJ).

### Quantification and statistical analysis

Raw microscopy images, deconvoluted Z-stacks and movies were transferred to FIJI (ImageJ) for final analysis and processing. Fluorescence traces were analysed in OriginPro 2021 (V. 9.8.0.200 [Academic]). Note that all fluorescence traces were generated from raw microscopy data and not deconvoluted data. Cell lengths were extracted from MicrobeJ (rod and cocci) ^55^ or by manual tracing in Fiji (filaments). Evaluations of statistical significance were performed using students T-tests in GraphPad Prism software (v.9.2). Levels of significance are indicated as follows: ns, not significant; *, *P* < 0.05; **, *P* < 0.01, ***, *P* < 0.001, ****, *P* < 0.0001

## Data availability

- All data reported in this paper is available within the manuscript or its supplementary material.
- This paper does not report original code.

## Acknowledgments

The authors want to thank Prof. Danielle Garsin (University of Texas) for the *E. faecalis* OG1RF strain and Prof. Scott Hultgren (University Washington in St. Louis) for the *K. pneumoniae* TOP52 strain. B.S. acknowledge support from the Australian Research Council through an ARC Future Fellowship (FT230100062). This study received funding from internal UTS and Australian Institute for Microbiology and Infection (AIMI) funding schemes to S.R. and B.S. The authors acknowledge the use of the Leica Stellaris confocal microscope in the Microbial Imaging Facility at AIMI in the Faculty of Science, University of Technology Sydney.

## Author contributions

B.S. conceptualised the study together with I.D.. C.A-K conduced the experiments together with B.S.. C.A-K and A.P constructed plasmids. I.D., C.A-K and B.S. analysed the data. S.A.R. and B.S. raised funding from UTS and AIMI for this project. C.A-K., A.P., I.D., and B.S contributed to the manuscript (writing and editing), all authors approved the final version.

